# Spectral projectors of temporally integrated functional brain networks

**DOI:** 10.64898/2026.03.20.707714

**Authors:** Vladi Sorkin, Yakir Menahem, Dmitry Patashov, Michal Balberg, Dmitry Goldstein

## Abstract

Functional-connectivity matrices mix pairwise association strength, spectral weighting, and eigenspace orientation, making it difficult to determine which components carry stable information about network organization. We develop a spectral-geometric framework that uses rank-*k* orthogonal projector matrices as subject-level representations of functional operators. The projector preserves eigenspace orientation in measurement coordinates while removing relative eigen-value weighting. We further compare two noncommuting temporal constructions—projecting a temporally integrated operator and averaging projectors estimated from local operators— and examine how the resulting representations depend on temporal scale and retained rank. We demonstrate the framework in resting-state fNIRS recordings from 99 four-month-old infants exposed to monolingual or bilingual language environments. Pearson correlation favored temporal integration before projection and exhibited a localized intermediate-rank regime at shorter scales, whereas multitaper coherence favored projection before integration and showed a broader regime at longer scales. Reverse engineering of the Pearson representation showed that the intermediate-rank regime was carried primarily by angular relations among channel embeddings, whereas a secondary near-full-rank regime corresponded to a distinct bottom-of-spectrum channel-occupancy carrier. A fully nested two-rank analysis repeatedly retained both regimes. These results show that temporally integrated functional-network organization depends jointly on operator definition, temporal support, projection order, and retained rank. The framework provides an interpretable way to separate complementary spectral carriers of network structure, with group labels used only for cross-fitted validation rather than for representation construction.

**Key Points:**

- Rank-*k* spectral projectors provide an interpretable, basis-invariant representation of functional-network eigenspace geometry in the original measurement coordinates.
- Temporal integration and spectral projection are noncommuting operations, and their relative informativeness depends on the functional operator and temporal scale.
- Distinct retained-rank regimes reveal complementary geometric carriers of functional-network organization, including relational dominant-subspace structure and bottom-of-spectrum channel occupancy.

## 1 Introduction

Functional connectivity (FC) describes statistical dependencies among distributed neural signals and is widely used to study brain organization (Bullmore and Sporns, 2009; Sporns, 2011; Fornito et al., 2016). Unlike structural connectivity, however, FC is constructed rather than directly observed: its organization depends on how interregional dependence is defined, estimated, and integrated over time. Different estimators can therefore yield materially different network representations (Friston, 2011; Bastos and Schoffelen, 2016; Liu et al., 2025). This raises a basic representational question: which properties of a functional operator reflect organized network structure, and which remain tied to spectral weighting or to weaker directions that may be unstable under finite-sample estimation?

Temporal support is part of the same problem. Conventional FC analyses often summarize an entire recording with a single matrix, while dynamic-FC approaches use local windows to characterize time-varying configurations (Hutchison et al., 2013; Allen et al., 2014; Calhoun et al., 2014; Zalesky et al., 2014; Lurie et al., 2020). Short windows provide temporally localized but variable estimates, and apparent window-to-window fluctuations can arise even under stationary or weak-signal conditions (Leonardi and Van De Ville, 2015; Hindriks et al., 2016). In the present framework, local windows are not interpreted as independent dynamic states. Instead, they serve as local estimators that are combined into a subject-level representation at a specified temporal scale. The relevant question is not only how much temporal support is used, but also how local operators are combined and whether temporal integration should precede or follow extraction of their spectral geometry.

A symmetric functional operator contains several conceptually distinct components. Its eigen-vectors define coordinated directions in signal space, while its eigenvalues determine the relative weighting of those directions. Entrywise analysis of the original operator consequently mixes association strength, spectral weighting, eigenspace orientation, and weaker finite-sample components. Individual eigenvectors are additionally sign-indeterminate and may rotate within nearly degenerate eigenspaces without a meaningful change in the underlying subspace (Davis and Kahan, 1970).

Spectral and subspace-based representations are established mathematical tools and have entered brain-network analysis in several forms. Structural connectome eigenmodes have been used as bases for representing functional activity, and joint subspace mappings have been used to relate structural and functional connectomes (Atasoy et al., 2016; Preti and Van De Ville, 2019; Ghosh et al., 2023). The methodological contribution does not lie in the orthogonal projector as a mathematical object. It lies in using the rank-*k* spectral projector of a functional operator as an interpretable subject-level matrix-valued representation in the original measurement coordinates, and in studying how this representation changes jointly with temporal support, retained rank, and the order of temporal integration and spectral projection.

For a real symmetric functional operator *C* ∈ R*^p^*^×^*^p^*, let *U_k_* ∈ R*^p^*^×^*^k^* contain the eigenvectors associated with its *k* largest eigenvalues. We define

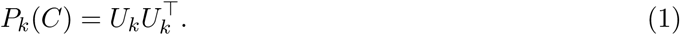

The projector depends only on the retained subspace and is invariant to eigenvector sign changes and to orthogonal rotations of the basis within that subspace. Unlike the weighted rank-*k* reconstruction *U_k_Λ_k_U^T^_k_*, it removes relative eigenvalue weighting while preserving how the retained geometry is embedded in the original measurement coordinates (Fig. 1A) (Edelman et al., 1998; Absil et al., 2008).

**Figure 1:**
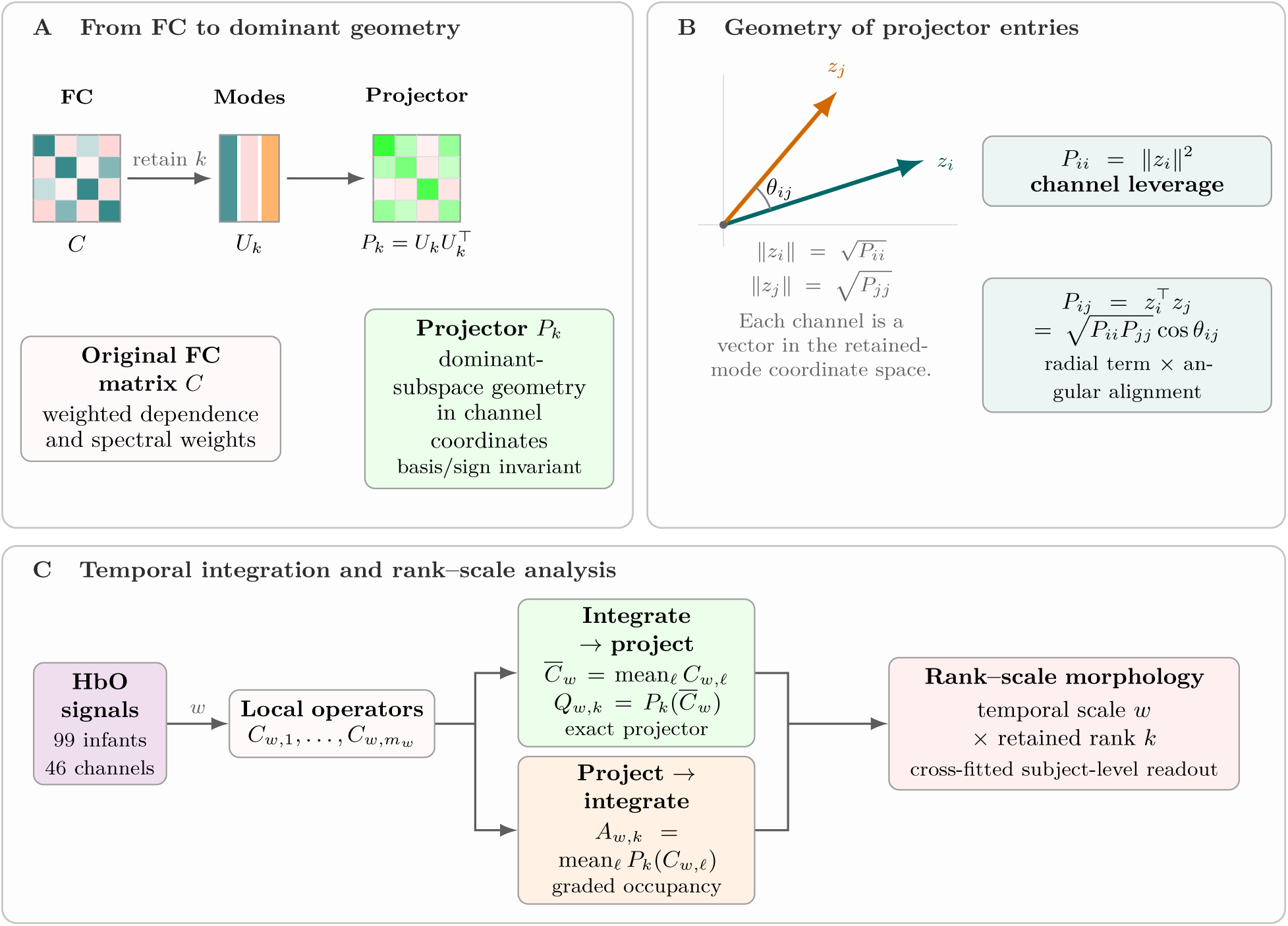
Projector-derived functional-network representation and analysis framework. **A**, a functional-connectivity operator *C* retains weighted statistical dependence, while *P_k_*= *U_k_U* ^T^ retains the basis-invariant geometry of its leading *k*-dimensional eigenspace in channel coordinates. **B**, the *i*th row *z*^T^ of *U_k_* embeds channel *i* across the retained modes. Diagonal entries quantify channel leverage; off-diagonal entries combine the leverage of two channels with their signed angular alignment. **C**, local Pearson-correlation or multitaper-coherence operators are combined either before or after spectral projection. The resulting matrices are examined across temporal scale *w* and retained rank *k*; group labels enter only through the cross-fitted subject-level readout.

**Figure 2:**
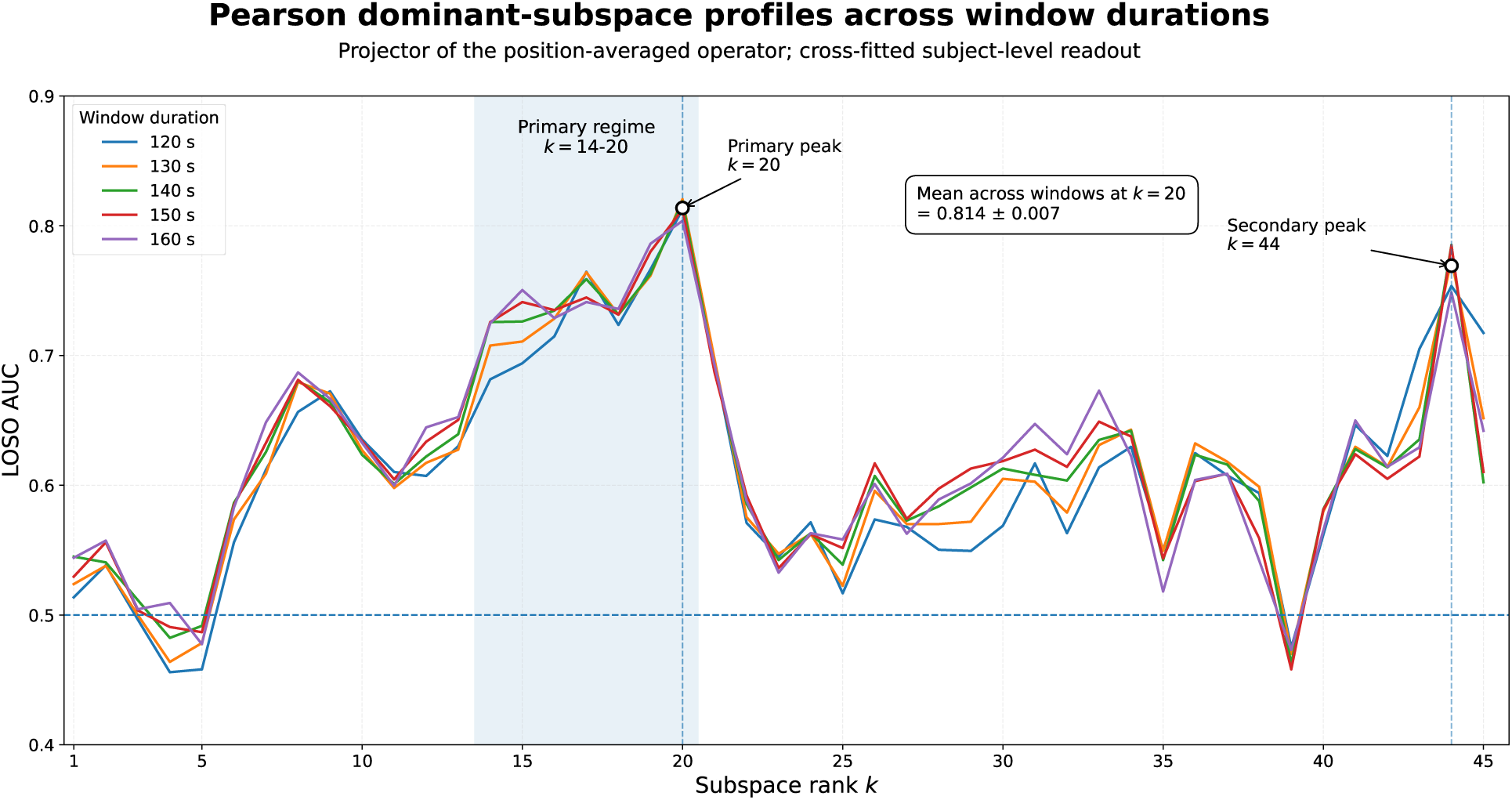
Pearson rank–scale morphology across the complete rank range. Outer leave-one-subject-out AUC across retained rank *k* = 1, …, 45 for window durations of 120–160 s. The shaded region marks the principal intermediate-rank regime (*k* 14–20), with *k* = 20 retained as the representative rank. The complete profile also reveals a secondary near-full-rank rise at *k* = 44, examined separately in the post hoc rank-dependent analysis.

The entries of *P_k_* have a direct geometric interpretation. Let *z_i_^T^ = e^T^_i_ U_k_* denote the *i*th row of *U_k_*, that is, the coordinates of channel *i* across the retained eigenmodes. Then

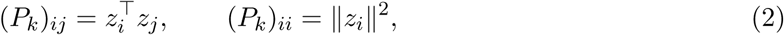

and, whenever both diagonal entries are nonzero,

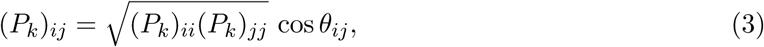

where *θ_ij_* is the angle between the retained-mode participation profiles *z_i_* and *z_j_* (Fig. 1B).

A diagonal entry measures channel leverage—the total participation of a channel in the retained subspace—while an off-diagonal entry combines the marginal participation of two channels with their signed angular alignment across retained modes. The normalized quantity

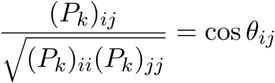

isolates relational angular geometry from leverage.

This interpretation is deliberately different from that of an FC matrix. An entry *C_ij_* represents an estimated statistical dependence between two measured signals. By contrast, an entry (*P_k_*)*_ij_* describes how two channels are jointly embedded in a retained eigenspace of the complete operator. The original FC matrix retains weighted statistical dependence, while the projector-derived matrix retains basis-invariant subspace geometry. The two representations are complementary rather than interchangeable.

Temporal integration and spectral projection define two noncommuting constructions (Fig. 1C).

Local functional operators may first be averaged and the integrated operator projected,

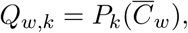

or each local operator may first be projected and the resulting projectors averaged,

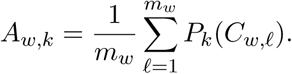

The former is an exact rank-*k* projector of the temporally integrated operator. The latter is generally not a projector; it is a graded subspace-occupancy matrix that summarizes geometry recurring across local estimates. Because spectral projection is nonlinear,

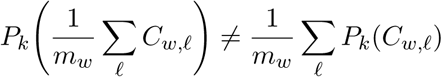

in general. The relative informativeness of the two constructions may depend on both the functional operator and temporal support.

Within a fixed operator family and temporal construction, the framework yields a two-axis family of matrix-valued representations. Window duration *w* controls temporal support, while retained rank *k* controls the dimension of the represented subspace. We treat temporal scale and rank as scientific coordinates, not solely as tuning parameters. Their joint variation defines a rank–scale morphology in which stable regimes and transitions may reveal changes not only in the amount of retained information but also in the geometric component through which that information is expressed.

We evaluate this framework using Pearson correlation and multitaper coherence as complementary time- and frequency-domain functional operators in a public resting-state fNIRS dataset comprising 99 four-month-old infants exposed to monolingual or bilingual language environments (Blanco et al., 2022). Resting-state fNIRS has been used to characterize large-scale functional organization during early development (Homae et al., 2011; Hu et al., 2020). Studies using fNIRS to examine bilingual brain organization remain comparatively limited, particularly in infancy (Pliatsikas, 2024; Blanco et al., 2025). We use this cohort as a challenging developmental-neuroimaging application in which group-associated variation provides an external readout of information retained by alternative network representations.

Previous analyses of this cohort found no corrected group differences in subject-specific temporal-ICA maps or in the expression of components derived from Pearson-connectivity matrices (Blanco et al., 2021). A later frontal-lobe analysis reported largely null differences in mean FC strength, together with differences in selected lateralization-related graph and directional measures (Gao et al., 2022). The present study addresses a different representational target: whole-montage subspace geometry of temporally integrated functional operators.

We ask three methodological questions. First, do projector-derived representations exhibit structured rank–scale organization beyond isolated high-readout configurations? Second, does the preferred order of temporal integration and spectral projection depend on the functional operator? Third, when distinct rank regimes emerge, which geometric components of the projector carry their label-associated information?

Functional operators and projector-derived matrices are constructed without group labels. Labels enter only through strictly cross-fitted subject-level readouts used to characterize label-associated variation retained by each representation. These readouts are not intended to establish diagnostic separability, causal effects of bilingual exposure, or a global difference in functional connectivity. The objective is to characterize functional-network representations and the methodological choices that make different components of their spectral geometry visible.

## 2 Methods

### 2.1 Study design and data

We developed a matrix-valued spectral framework in which eigenspaces of symmetric functional-connectivity operators are represented directly by orthogonal projector matrices in measurement coordinates. The framework compares two noncommuting orders of temporal integration and spectral projection and characterizes the resulting representations across temporal scale and retained rank. Pearson correlation and multitaper coherence were used as complementary time- and frequency-domain functional operators.

We analyzed the open RS4 task-free fNIRS dataset (Blanco et al., 2022). The original sample comprised 104 healthy, full-term, four-month-old infants recorded during natural sleep. Following the exclusions used in the original group analysis (Blanco et al., 2021), 99 infants remained: 36 Basque–Spanish bilingual, 30 Spanish monolingual, and 33 Basque monolingual. The two monolingual subgroups were combined, yielding 63 monolingual and 36 bilingual infants.

Data were acquired with a NIRx NIRScout system at *f_s_* = 8.93 Hz using 760- and 850-nm wavelengths. We used the distributed, analysis-ready, global-signal-regressed HbO data, rsData.GSR_oxy. The original preprocessing included removal of poor-quality recording segments, wavelet motion correction, conversion to hemoglobin concentration changes, temporal nuisance regression, and cropping to 5000 samples (≈ 560 s). After the dataset-provided channel selection and reordering,

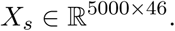

The channel selection was treated as already incorporated and was not applied again. The infant was the statistical unit. Temporal windows were used only to construct within-subject operators and were never treated as independent observations.

### Ethics

The present study is a secondary analysis of publicly available, de-identified data. The original data collection was approved by the local ethics committee at the Basque Center on Cognition, Brain and Language (Donostia–San Sebastián, Spain). Parents were informed about the study procedures, their right to withdraw, and the planned sharing of non-identifiable data, and written informed consent was obtained from all parents before data acquisition (Blanco et al., 2022).

### 2.2 Centered temporal-window sampling

For window duration *w*, the window length was *L_w_* = round(*wf_s_*). Window positions were sampled with a requested 10-s stride, Δ = round(10*f_s_*), and centered within the retained recording. The number of admissible windows and their zero-based start positions were

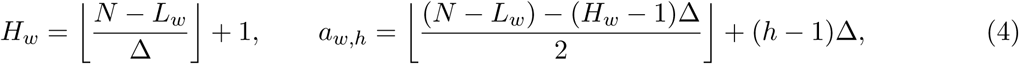

for *h* = 1*, …, H_w_*. This rule imposed the same endpoint-neutral positional sampling on all subjects at a given temporal scale.

### 2.3 Projector-derived functional-network representations

For a real symmetric functional operator *C* = *V* Λ*V* ^T^, with eigenvalues ordered from largest to smallest, let *U_k_* = [*v*_1_*, …, v_k_*]. The retained eigenspace was represented by

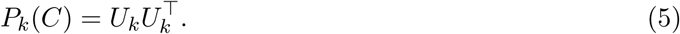

*P_k_*(*C*) is an orthogonal rank-*k* projector and is invariant to eigenvector sign changes and to orthogonal changes of basis within the retained subspace.

For local operators *C^o^_s,h_(w)*, we compared integration before projection,

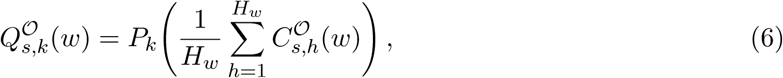

with projection before integration,

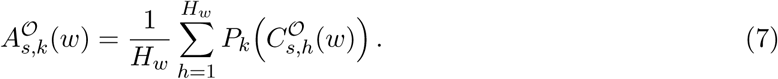

Here, *Q* is an exact rank-*k* orthogonal projector of the temporally integrated operator. In contrast, *A* is a positive-semidefinite, generally non-idempotent subspace-occupancy matrix satisfying 0 ⪯ *A* ⪯ *I* and tr(*A*) = *k*.

Projector-derived matrices were vectorized using their upper triangles including the diagonal, giving

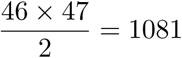

coordinates. For operator baselines with a constant diagonal, only the strict upper triangle was retained.

### 2.4 Pearson correlation operators

For each subject, temporal scale, and window position, we computed

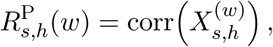

followed by shrinkage

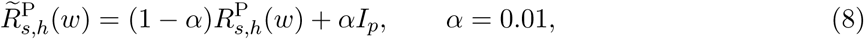

with unit diagonal enforced after regularization.

The primary Pearson representation integrated local operators before projection,

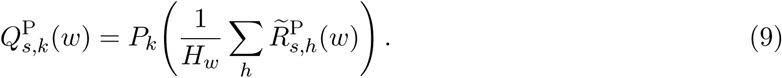

Rank–scale morphology was evaluated for

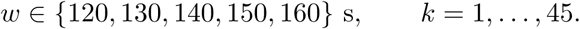

Rank 46 was excluded because *P*_46_ = *I_p_*for every subject. The fixed follow-up analyses used *w* = 140 s and *k* = 20, selected as the representative Pearson configuration from a contiguous intermediate-rank region in the descriptive rank–scale surface, not from an isolated numerical maximum.

At this configuration, the complementary construction was

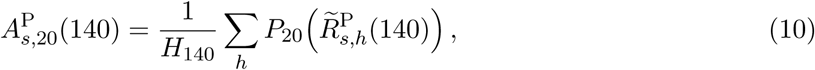

and a secondary control reprojected this matrix,

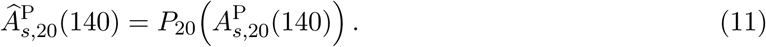

Label-free differences between *Q* and *A* were summarized by idempotence error ∥*A*^2^ − *A*∥*_F_*, normalized Frobenius distance *‖Q−A‖_F_/2k* and normalized overlap tr(*QA*)*/k*.

Positional-sampling robustness was evaluated at the representative configuration using requested strides of 5, 10, 15, 20, 30, 40, and 50 s (Supporting Table S1).

### 2.5 Multitaper coherence operators

Local windows were demeaned channel-wise and analyzed using multitaper spectral estimation with time–bandwidth product *NW* = 3, *K* = 5 DPSS tapers, and equal taper weights (Thomson, 1982; Babadi and Brown, 2014). For tapered Fourier coefficients *Y_s,h,i,m_*(*f*),

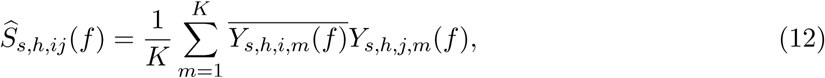

and magnitude-squared coherence was

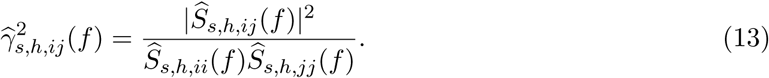

The local coherence operator was obtained by uniformly averaging over Fourier frequencies in 0.03–0.08 Hz,

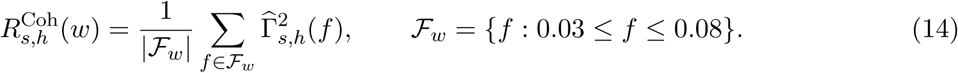

Matrices were symmetrized and assigned unit diagonal.

The primary coherence representation projected each local operator before temporal integration,

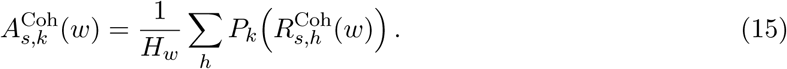

A preliminary temporal-scale scan used fixed rank *k* = 20 and windows from 130 to 300 s. The focused rank–scale analysis used

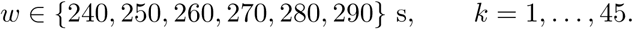

The fixed follow-up analyses used *w* = 260 s and *k* = 20, selected as the representative coherence configuration from a broad region in the descriptive rank–scale surface. The complementary integration-before-projection construction,

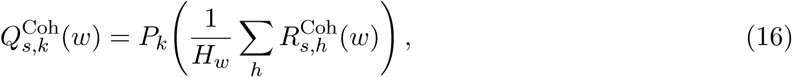

was evaluated over the same focused grid.

Position-averaged raw coherence and channel-wise log band power served as frequency-domain controls. Sensitivity to taper number was examined for *K* = 2, 3, 4, 5 DPSS tapers at the representative configuration. An additional focused temporal-scale sensitivity analysis compared *K* = 3, 4, 5 across window durations of 250–280 s at fixed rank *k* = 20. Additional results are reported in Supporting Figure S3 and Supporting Table S4.

### 2.6 Rank–scale characterization

Temporal scale and retained rank were treated as scientific coordinates of the representation, not solely as hyperparameters. Rank–scale surfaces were characterized by the location, width, and persistence of elevated readout regions, with isolated numerical maxima not used as the primary basis for interpretation.

For Pearson correlation, the principal representative configuration was defined from a stable contiguous region of the descriptive rank–scale surface instead of the single highest-AUC cell. A secondary feature identified in the complete rank profile was examined separately and did not redefine the principal configuration. Its geometric interpretation and the subsequent two-rank analysis were explicitly designated as post hoc or exploratory.

### 2.7 Cross-fitted subject-level readout and statistical inference

Fixed representations were evaluated using strict outer leave-one-subject-out cross-validation. In each of 99 outer folds, one infant was held out and all data-dependent transformations were estimated using only the remaining 98 infants (Varoquaux et al., 2017; Scheinost et al., 2019).

Within each training fold, coordinates with nonfinite or near-zero variance were removed and the remaining features standardized. Partial least squares regression with five latent components was fitted to the standardized training features and labels, followed by *ℓ*_2_-regularized logistic regression with *C* = 0.5 and inverse-frequency class weighting. The held-out subject was transformed using only parameters estimated from the corresponding training fold.

Group labels were not used to define windows, estimate functional operators, construct eigenspaces, or form projector-derived matrices. Labels entered only through the fold-specific PLS and logistic-regression stages.

Outer-fold probabilities were summarized by area under the receiver-operating-characteristic curve (AUC). Balanced accuracy, monolingual specificity, and bilingual sensitivity at a probability threshold of 0.5 were secondary summaries. These quantities were treated as cross-fitted validation readouts and not as estimates of diagnostic utility.

For fixed representations, uncertainty in AUC and paired AUC differences was estimated using stratified subject bootstrap resampling of saved outer-LOSO predictions, with 5000 replicates unless otherwise stated. The matched rank-dependent decomposition and nested rank-pair analyses used 10,000 bootstrap replicates.

The fixed principal Pearson configuration was additionally evaluated using 9,999 global label permutations. For each permutation, PLS and logistic regression were refitted within every outer training fold. The one-sided Monte Carlo *p*-value used the finite-sampling correction of Phipson and Smyth (2010). This test was conditional on the fixed representation and readout specification and did not correct for the preceding descriptive rank–scale exploration.

Robustness to unequal group size was examined using 50 repeated balanced cohorts containing all 36 bilingual infants and 36 monolingual infants. The complete outer-LOSO pipeline was rerun at the fixed Pearson configuration; additional aggregation details are reported in Supporting Table S2.

### 2.8 Fully nested adaptive rank selection

Adaptive rank selection was evaluated at the fixed Pearson temporal scale *w* = 140 s using projectors *P_s,k_* for all *k* = 1*, …,* 45. Outer validation remained leave-one-subject-out. Within each outer training set, candidate ranks were evaluated using a common stratified nine-fold inner cross-validation. The complete training-only variance filtering, standardization, five-component PLS, and balanced *ℓ*_2_-regularized logistic-regression pipeline was refitted within every inner fold.

For single-rank selection, inner out-of-fold predictions were generated for each candidate *k*, and the rank maximizing pooled inner-OOF AUC was selected. Exact ties were resolved in favor of the smaller rank. The selected representation was refitted on the complete outer training set and evaluated once on the held-out infant.

Inspection of the descriptive Pearson rank profile motivated an additional exploratory analysis allowing two rank-specific branches to be retained simultaneously. The decision to examine a two-branch representation was made after inspection of the rank profile; selection and evaluation of the individual ranks nevertheless remained fully nested.

For each candidate rank *k*, let *ℓ_i_*(*k*) denote its inner-OOF logistic decision score for an outer-training subject. These scores were standardized using only the outer-training inner-OOF distribution,

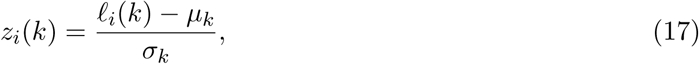

where *µ_k_* and *σ_k_* are the mean and standard deviation of the inner-OOF decision scores for rank *k*. Every distinct pair *k_a_ < k_b_* was evaluated using fixed equal-weight late fusion,

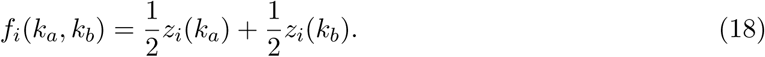

The pair maximizing pooled inner-OOF AUC was selected. Exact ties were resolved lexicographically, favoring the smaller *k_a_* and then the smaller *k_b_*. Neither the fusion weight nor the decision threshold was optimized.

The two selected rank-specific readouts were refitted independently on the complete outer training set. Each held-out decision score was standardized using the corresponding training-only inner-OOF mean and standard deviation, and the two standardized scores were averaged with the same fixed weights. The held-out infant and its label could not affect rank selection, score normalization, or fusion.

The primary adaptive comparison was the fully nested two-rank equal-fusion procedure against the fully nested single-rank selector. Allowing two ranks was scientifically motivated by the descriptive profile and was therefore exploratory. The reported outer-LOSO performance nevertheless evaluates the specified adaptive procedures without outer-test leakage. Selection-frequency diagnostics are reported in the Supporting Information.

### 2.9 Geometric decomposition of projector representations

At *w* = 140 s and *k* = 20, let

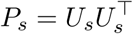

and let *z^T^_s,i_* denote row *i* of *U_s_*. Then

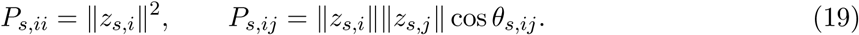

Diagonal entries were treated as channel leverage. With a 10^−12^ numerical floor applied to diagonal terms, pairwise radial and normalized angular components were

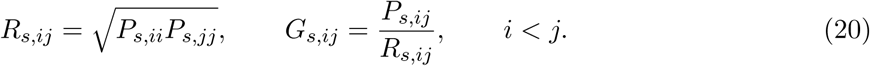

Five fixed feature families were evaluated using the same outer-LOSO pipeline: diagonal leverage, radial terms, normalized angular relations, raw off-diagonal entries, and the complete upper triangle.

For exploratory spatial description, angular group effects were summarized using Hedges’ *g*. Channel-level relational involvement was

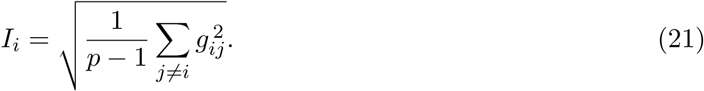

Spatial summaries were interpreted only in measurement-channel coordinates.

### 2.10 Post hoc analysis of rank-dependent representational carriers

A secondary near-full-rank feature identified during descriptive inspection of the complete Pearson rank profile was examined in a separate post hoc analysis. This analysis did not redefine the principal Pearson configuration.

For the 46-channel integrated Pearson operator at *w* = 140 s, this feature corresponded to

*k* = 44, for which

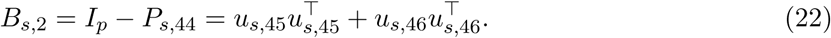

The near-full-rank *P_s,_*_44_ representation is therefore equivalently determined by the invariant two-dimensional bottom-spectrum projector *B_s,_*_2_. Individual bottom eigenvectors were not interpreted.

For the complete matrix-valued representation, *P_s,_*_44_ and *B_s,_*_2_ are related by the fixed affine bijection

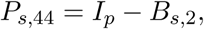

so they contain the same between-subject information. We used *B_s,_*_2_ for reverse engineering because it gives a lower-dimensional and more direct representation of the corresponding bottom-spectrum carrier.

This equivalence applies to the complete matrix-valued representation. The nonlinear radial and angular decompositions of *B_s,_*_2_, however, are not algebraically identical to decompositions of *P_s,_*_44_. They were used specifically to characterize the geometry of the complementary bottom-tail representation.

To test whether the intermediate- and near-full-rank representations were supported by the same geometric components, *P_s,_*_20_ and *B_s,_*_2_ were examined using matched decompositions. For *B_s,_*_2_, diagonal bottom-tail occupancy was

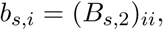

and the corresponding radial and normalized angular terms were

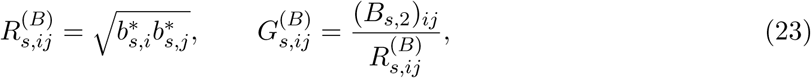

where *b^*^_s,i_ = max(b_s,i_, 10^−12^).*

The same five feature families were evaluated for both rank regimes: diagonal entries, radial terms, normalized angular relations, raw off-diagonal entries, and the complete upper triangle. All were evaluated with the same strict outer-LOSO readout. The purpose of this analysis was to characterize the geometric carrier of each fixed rank regime, not to select a new representation.

For spatial description, *P*_20_ was summarized using angular relational involvement, while *B*_2_ was summarized using Hedges’ *g* for channel-wise diagonal occupancy. Signed effects were defined as bilingual minus monolingual. These maps were exploratory and were not interpreted as cortical localization.

### 2.11 Exploratory cross-operator complementarity

Complementarity between the representative Pearson and coherence branches was explored using their saved outer-LOSO decision scores. The fixed fusion was an untrained equal-weight mean in logit space,

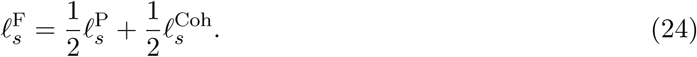

No meta-classifier or learned fusion weight was used. Pearson and Spearman score correlations, prediction disagreement, and paired bootstrap comparisons were treated as descriptive measures of complementarity.

## 3 Results

Projector-derived functional-network representations showed structured, operator-dependent variation across temporal scale *w*, retained rank *k*, and the order of temporal integration and spectral projection. We first characterize the Pearson and coherence rank–scale morphologies, then compare the two temporal constructions, identify the geometric carriers of distinct Pearson rank regimes, and finally evaluate adaptive rank selection.

### 3.1 Pearson correlation shows a localized intermediate-rank regime

For the projector of the position-averaged Pearson operator, window durations of 120–160 s showed a consistent elevated region at approximately *k* = 14–20 (Fig. 2). The readout increased toward intermediate rank and declined after *k* = 20, with similar morphology across all five temporal scales. The complete Pearson profile also showed a secondary near-full-rank rise around *k* = 44, examined separately below.

At *k* = 20, mean AUC across the five window durations was 0.814 (SD = 0.007). We retained *w* = 140 s and *k* = 20 as the representative Pearson configuration. Its outer-LOSO readout yielded AUC = 0.8188, balanced accuracy = 0.7698, monolingual specificity = 0.7619, and bilingual sensitivity = 0.7778.

The result was robust to positional sampling: requested strides from 5 to 50 s yielded AUCs of 0.7910–0.8192. Repeated equal-sized cohorts preserved the pattern, with a mean AUC of 0.7557 across 50 balanced cohorts and an AUC of 0.7945 after aggregation of repeated cross-fitted predictions (Supporting Tables S1–S2).

At the fixed representative configuration, 9,999 global label permutations gave a one-sided Monte Carlo *p* = 0.0002. This inference was conditional on the fixed representation and readout specification and did not correct for the preceding descriptive rank–scale exploration (Supporting Table S3).

Exploratory controls using Spearman and Gaussian-copula dependence retained rank-dependent organization, while a heat-kernel construction produced a different rank profile (Supporting Figures S4–S5). Rank-dependent structure was therefore not specific to Pearson correlation, although its spectral location depended on the operator.

### 3.2 Coherence shows a broader regime at longer temporal scales

Multitaper coherence showed a broader elevated region at longer temporal scales. For the mean-local-projector construction, elevated readout extended approximately across *k* = 17–24 and *w* = 250–280 s (Fig. 3).

**Figure 3:**
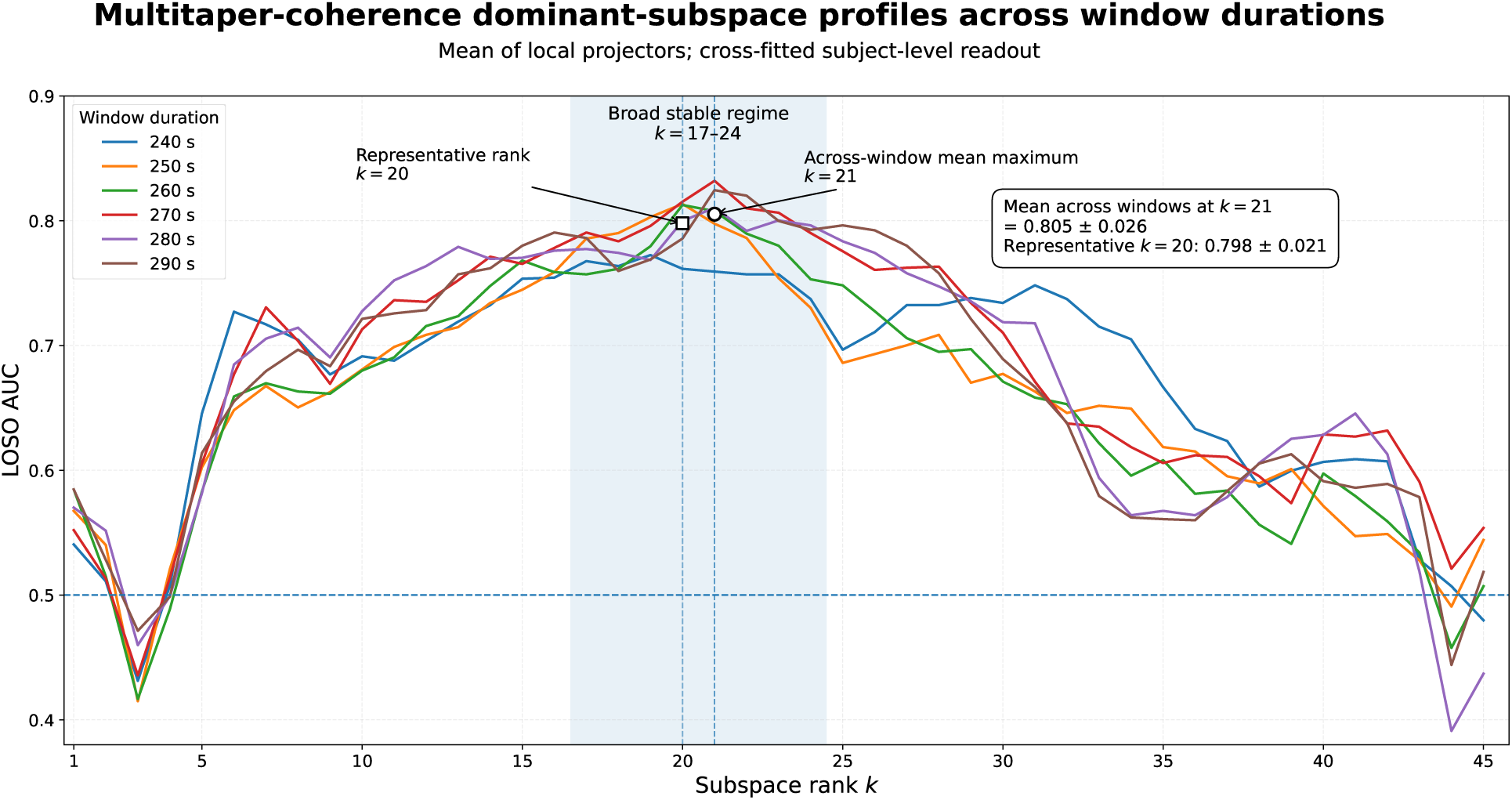
Coherence rank–scale morphology across the complete rank range. Outer leave-one-subject-out AUC for the multitaper-coherence mean-local-projector representation across retained rank *k* = 1*, …,* 45 and window durations of 240–290 s. The elevated region spans approximately *k* = 17–24. The representative configuration was *w* = 260 s and *k* = 20.

**Figure 4:**
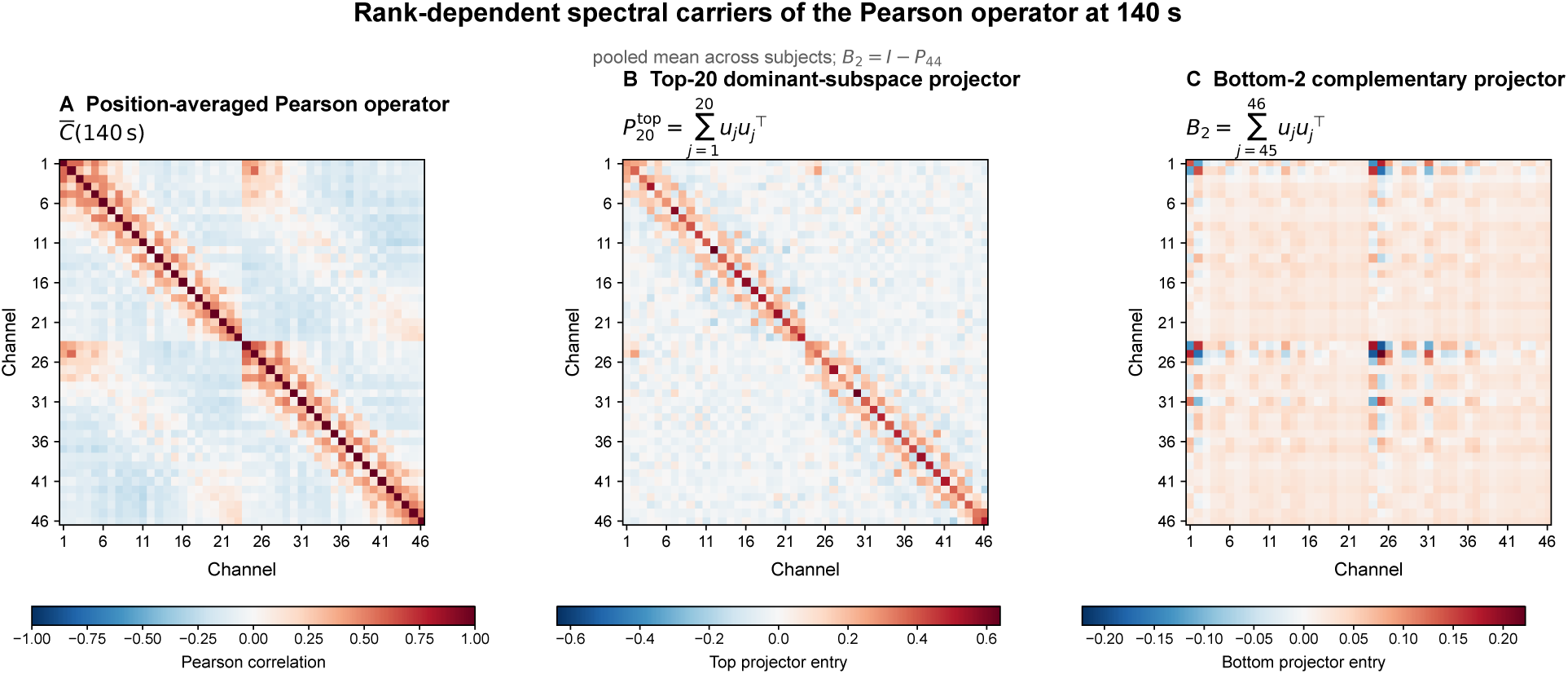
Rank-dependent spectral carriers of the temporally integrated Pearson operator at 140 s. For visualization, the position-averaged Pearson operators were pooled across all subjects before eigendecomposition; no group labels entered this construction. **A,** pooled position-averaged Pearson operator *C^-^*(140 s). **B**, rank-20 dominant-subspace projector *P_20_ = Σ^20^_j=1_ u_j_u^T^_j_*, illustrating the intermediate-rank spectral representation. **C**, complementary bottom-two projector *B_2_ = Σ^46^_j=45_ u_j_u^T^_j_ = I − P_44_*, which provides the low-dimensional representation associated with the near-full-rank *k* = 44 regime. Color scales are panel specific and are intended to reveal the structure of each representation rather than to support direct comparison of entry magnitudes across panels.

The largest value in the descriptive grid occurred at *w* = 270 s, *k* = 21 (AUC = 0.8320), but this isolated cell was not treated as a validated optimum. The representative *w* = 260 s, *k* = 20 configuration yielded AUC = 0.8126 and balanced accuracy = 0.7004.

Controls without dominant-subspace projection were substantially weaker: position-averaged raw coherence reached approximately 0.62 AUC, and channel-wise band power remained near 0.60– 0.61. The representative coherence readout remained informative for three to five DPSS tapers across neighboring window durations at fixed *k* = 20, with *K* = 5 providing the most consistently high AUC (Supporting Figure S3 and Supporting Table S4). No standalone label-permutation test was performed for coherence, so these values are reported as descriptive cross-fitted readouts.

Pearson and coherence therefore exhibited distinct rank–scale morphologies: a localized intermediate-rank Pearson regime at shorter windows and a broader coherence regime at longer windows.

### 3.3 The preferred order of temporal integration and spectral projection is operator dependent

The two operator families favored opposite orders of temporal integration and spectral projection.

At the representative Pearson configuration, projecting the position-averaged operator yielded AUC = 0.8188, whereas averaging local rank-20 projectors yielded AUC = 0.5970. The paired difference was ΔAUC = 0.2218 (95% bootstrap interval [0.1234, 0.3254]). Reprojecting the mean-local-projector matrix did not recover the stronger readout (AUC = 0.5670).

The two Pearson constructions were also geometrically distinct without reference to labels. The mean-local-projector matrix had mean idempotence error 1.090, normalized Frobenius distance from the projector of the integrated operator 0.322, and normalized overlap tr(*QA*)*/k* = 0.734.

Coherence showed the reverse ordering. Averaging local projectors at *w* = 260 s, *k* = 20 yielded AUC = 0.8126, whereas projecting the position-averaged coherence operator yielded AUC = 0.7002. The paired difference was ΔAUC = 0.1124 (95% bootstrap interval [0.0322, 0.1953]).

Temporal integration and spectral projection were not interchangeable: integration before projection was favored for Pearson, whereas projection before integration was favored for coherence (Supporting Tables S5–S6).

### 3.4 Relational angular geometry carries the principal Pearson regime

Figure 4 provides a label-free overview of the integrated Pearson operator and the intermediate- and near-full-rank spectral carriers analyzed below.

At *w* = 140 s and *k* = 20, the Pearson projector was decomposed into diagonal channel leverage, pairwise radial terms, normalized angular relations, raw off-diagonal entries, and the complete upper triangle (Fig. 5A).

**Figure 5:**
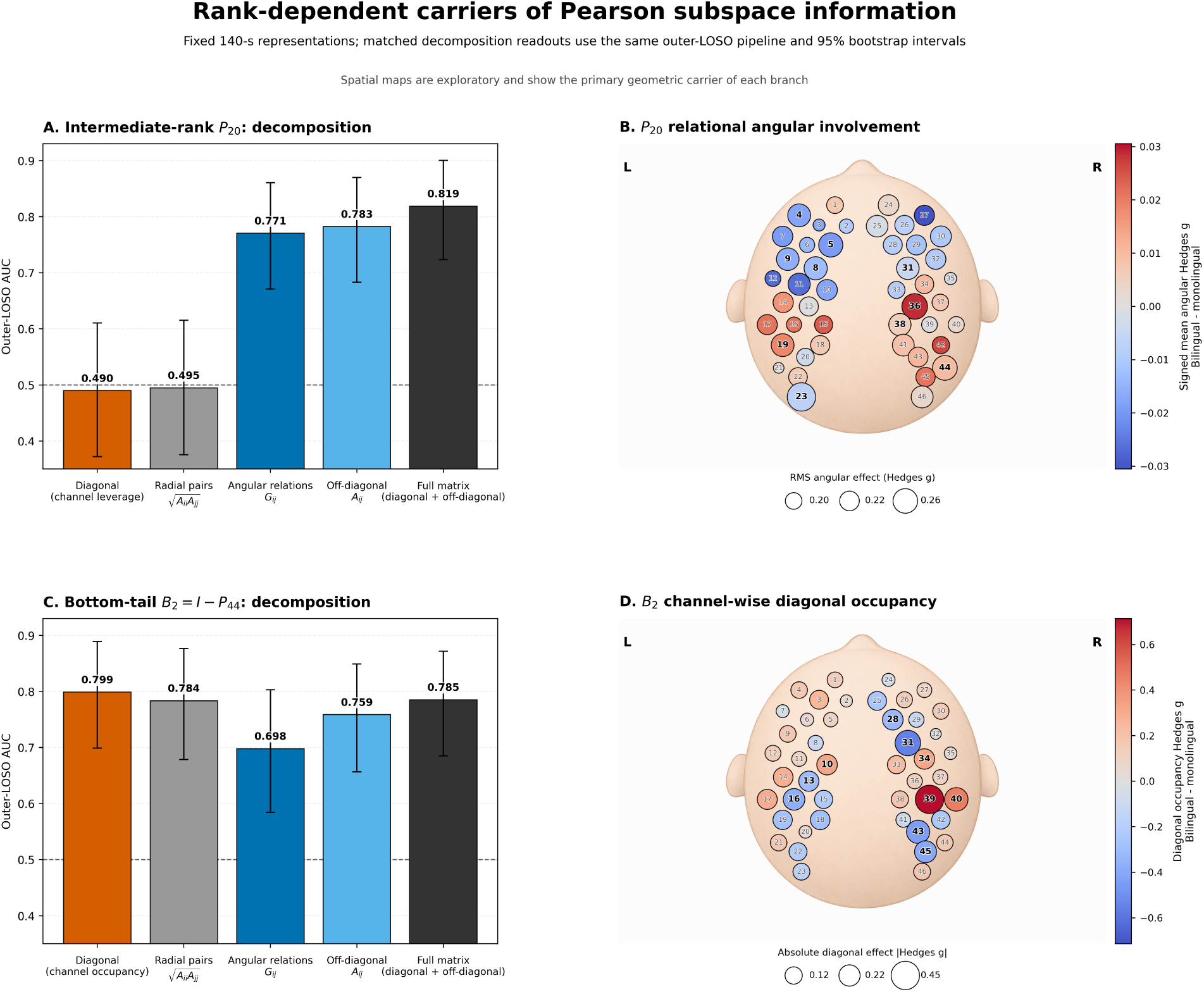
Rank-dependent representational carriers of Pearson subspace information. **A,** matched decomposition of the principal intermediate-rank projector *P_20_*. **B**, exploratory channel-level angular involvement for *P_20_*. **C**, matched decomposition of the complementary bottom-tail projector *B_2_ = I − P_44_*. **D**, exploratory channel-wise differences in diagonal bottom-tail occupancy. Error bars in A and C denote 95% stratified-bootstrap intervals. Spatial panels are descriptive measurement-space summaries and are not interpreted as independently confirmed cortical localization.

Diagonal leverage and radial terms were at chance (AUC = 0.490 and 0.495, respectively). In contrast, normalized angular relations yielded AUC = 0.771, raw off-diagonal entries 0.783, and the complete projector 0.819 (Supporting Table S7).

The principal intermediate-rank Pearson readout was carried mainly by relations among channel embeddings, with little contribution from marginal channel leverage. The corresponding angular involvement was distributed across the measurement montage, with no single channel dominating the pattern (Fig. 5B). These spatial quantities were treated as exploratory measurement-space summaries.

### 3.5 Post hoc analysis reveals distinct rank-dependent representational carriers

The secondary near-full-rank Pearson regime around *k* = 44, identified during inspection of the complete rank profile, was examined explicitly post hoc. For the 46-channel integrated operator, the rank-44 projector is equivalently determined by the two-dimensional complement

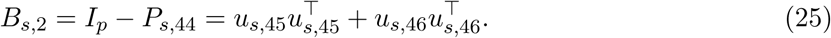

Figure 4 illustrates the corresponding spectral organization in the label-free pooled Pearson operator. The intermediate-rank *P*_20_ representation occupies the dominant portion of the spectrum, while the near-full-rank *P*_44_ regime has the compact complementary representation *B*_2_ at the bottom of the spectrum.

We analyzed the invariant bottom-tail projector *B*_2_, rather than individual bottom eigenvectors, using the same five feature families as for *P*_20_ (Fig. 5C).

The carrier pattern differed qualitatively. Diagonal bottom-tail occupancy yielded AUC = 0.799, radial terms 0.784, normalized angular relations 0.698, raw off-diagonal entries 0.759, and the complete *B*_2_ matrix 0.785.

Whereas the *P*_20_ regime was predominantly relational, the bottom-tail regime was expressed most strongly through channel-wise occupancy of the complementary subspace. The corresponding occupancy differences were distributed across the montage (Fig. 5D).

The matched decomposition showed that retained rank changes not only the strength of the readout but also its geometric carrier: relational angular organization at intermediate rank and channel-wise bottom-tail occupancy near full rank.

### 3.6 Fully nested two-rank selection recovers complementary Pearson regimes

The coexistence of the intermediate- and near-full-rank Pearson regimes motivated an exploratory test of whether allowing two rank-specific branches would preserve information that adaptive single-rank selection could not. Although the two-rank hypothesis was motivated after inspection of the rank profile, rank selection and outer evaluation were fully nested as described in Methods.

A strict nested single-rank selector over *k* = 1*, …,* 45 yielded AUC = 0.7465 and balanced accuracy = 0.7123, compared with 0.8188 and 0.7698, respectively, for the fixed principal *k* = 20 representation. The paired AUC difference was −0.0723 (95% bootstrap interval [−0.1455, −0.0057]). The inner procedure selected *k* = 20 in 72 of 99 outer folds (72.7%), *k* = 44 in 25 folds (25.3%), and *k* = 14 and *k* = 17 once each. Unrestricted single-rank selection therefore alternated primarily between the two Pearson rank regimes.

Allowing two ranks produced a substantially different result. The fully nested two-rank procedure yielded AUC = 0.8765 (95% bootstrap interval [0.7862, 0.9506]) and balanced accuracy = 0.7917. Its AUC exceeded that of nested single-rank selection by 0.1301 (95% paired-bootstrap interval [0.0551, 0.2156], two-sided bootstrap *p* = 0.0004).

The selected ranks were strikingly stable. The pair *k* = 20, 44 was selected in 97 of 99 outer folds (98.0%), and *k* = 19, 44 in the remaining two. Rank 44 was included in every outer fold, while an intermediate-rank branch was retained in every fold. The adaptive procedure thus consistently combined one intermediate-rank and one near-full-rank branch across the outer validation.

Relative to the fixed principal *k* = 20 representation, the nested two-rank readout was higher by 0.0578 AUC, although the paired interval included zero ([−0.0018, 0.1235], *p* = 0.0594). The strongest conclusion is not that adaptive two-rank selection outperformed the fixed principal representation. Instead, it substantially outperformed adaptive single-rank selection and, under fully nested rank selection, repeatedly recovered the same two regimes suggested by the descriptive morphology and characterized by the geometric decomposition.

Selection-frequency and inner-pair diagnostics are reported in the Supporting Information.

The principal, control, mechanistic, and adaptive cross-fitted readouts are summarized in Table 1.

**Table 1:** Summary of principal, control, mechanistic, and adaptive cross-fitted readouts. AUC and balanced accuracy (BA) summarize subject-level outer-LOSO readouts. The two-rank model was motivated after inspection of the rank morphology, but rank-pair selection, score normalization, and held-out evaluation were fully nested within the outer cross-validation.

| Operator / analysis | Representation | Scale / rank | Role | AUC | BA |
| --- | --- | --- | --- | --- | --- |
| Pearson | integrate → project | 140 s, $k = 20$ | Principal | 0.8188 | 0.7698 |
| Coherence | project → integrate | 260 s, $k = 20$ | Principal | 0.8126 | 0.7004 |
| Pearson | project → integrate | 140 s, $k = 20$ | Order control | 0.5970 | 0.6052 |
| Coherence | integrate → project | 260 s, $k = 20$ | Order control | 0.7002 | 0.6488 |
| Pearson | nested single-rank selection | 140 s, $k \in [1, 45]$ | Nested adaptive | 0.7465 | 0.7123 |
| Pearson | nested two-rank equal fusion | 140 s,<br>$k_a < k_b \in [1, 45]$ | Nested adaptive | 0.8765 | 0.7917 |
| Pearson | $P_{20}$: normalized angular relations | 140 s, $k = 20$ | Mechanistic | 0.7707 | 0.6984 |
| Pearson | $P_{44}$: complete near-full-rank projector | 140 s, $k = 44$ | Post hoc regime | 0.7853 | 0.6944 |
| Pearson | $B_2 = I - P_{44}$: diagonal occupancy | 140 s | Post hoc mechanistic | 0.7989 | 0.7361 |
| Pearson + coherence | fixed equal-logit fusion | 140 / 260 s | Exploratory | 0.8911 | 0.8175 |

### 3.7 Pearson and coherence preserve partly complementary geometry

Pearson and coherence readout scores were moderately related (Pearson *r* = 0.337, *p* = 0.00064; Spearman *ρ* = 0.307, *p* = 0.0020) and produced different binary predictions for 41.4% of infants.

An exploratory fixed equal-weight mean of the two outer-LOSO logits yielded AUC = 0.8911 and balanced accuracy = 0.8175. The paired AUC increase over coherence was 0.0785 (95% interval [0.0194, 0.1429], *p* = 0.0084), while the increase over Pearson was 0.0723 (95% interval [−0.0013, 0.1477], *p* = 0.0540).

The fusion was interpreted as exploratory evidence that time-domain Pearson and frequency-domain coherence preserve partly distinct subject-level geometry. It was not treated as a separately optimized predictive model.

## 4 Discussion

This study develops a matrix-valued spectral framework for representing the geometry of temporally integrated functional brain networks using orthogonal projector matrices. The central result is a structured dependence of the representation on temporal support, retained rank, operator definition, and the order of temporal integration and spectral projection. Pearson correlation showed a localized intermediate-rank regime at shorter temporal scales, whereas multitaper coherence showed a broader regime at longer scales. The complete Pearson rank profile also revealed a secondary near-full-rank regime supported by a geometrically distinct component of the same integrated operator. Retained rank and temporal scale therefore affect not only how strongly label-associated variation is expressed, but also the form of network geometry through which that variation becomes visible.

Orthogonal projectors are useful in this setting because they represent an eigenspace without depending on eigenvector signs or on the particular orthogonal basis chosen within that subspace. They preserve how the retained spectral geometry is embedded in the original measurement coordinates while removing relative eigenvalue weighting. The resulting representation is therefore distinct from a weighted functional-connectivity matrix and from an abstract subspace description alone: its matrix entries remain directly linked to channel participation and to relations among channel embeddings. The methodological contribution lies in using this projector-derived representation jointly across temporal scale and retained rank, and in examining how its geometry changes under different orders of temporal integration and spectral projection.

A central finding was the operator-dependent reversal in the preferred order of these two operations. Pearson correlation favored integration before projection, whereas coherence favored projection of local operators followed by temporal averaging. Because spectral projection is nonlinear,

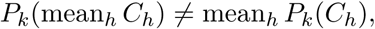

the two constructions are not expected to preserve identical geometry. The empirical results show that this noncommutativity has substantial representational consequences. One possible explanation is that averaging Pearson operators stabilizes distributed correlation structure before spectral truncation, while averaging local coherence projectors preserves spectral subspaces that recur across local frequency-domain estimates and may be attenuated when coherence matrices are averaged first. This interpretation remains mechanistic and is not uniquely established. Pearson and coherence also differ in frequency specificity, estimation variance, and preferred temporal scale. The supported conclusion is therefore that the preferred construction depends on the functional operator; the present data do not isolate operation order as the sole source of the difference between the two branches.

The rank–scale morphology similarly argues against treating retained rank as an ordinary tuning parameter. The principal Pearson regime occupied a contiguous region around *k* = 14–20 across neighboring temporal scales, while coherence showed a broader intermediate-rank region. These profiles describe how the represented network geometry changes as spectral directions are added. Rank can consequently be viewed as a coordinate of the representation: varying *k* changes which part of the operator’s spectral organization is retained and can expose different geometric properties of the same underlying functional operator.

The complete Pearson profile also demonstrated that informative structure was not confined to the dominant part of the spectrum. After the principal intermediate-rank regime declined, a secondary rise appeared near *k* = 44. For *p* = 46,

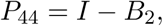

where *B*_2_ is the projector onto the complementary two-dimensional bottom-spectrum subspace. The near-full-rank representation can therefore be expressed through a compact low-dimensional complement. We analyzed the invariant projector *B*_2_, not individual bottom eigenvectors, because the latter remain sign-indeterminate and may rotate within their subspace. The appearance of this regime shows that bottom-spectrum directions can retain structured between-subject information despite their low spectral position.

Matched decomposition of *P*_20_ and *B*_2_ revealed qualitatively different representational carriers. For *P*_20_, diagonal channel leverage and pairwise radial terms were essentially uninformative in isolation, whereas normalized angular relations and raw off-diagonal entries retained most of the complete-projector readout. The principal intermediate-rank regime was therefore predominantly relational: label-associated variation was expressed in the arrangement of channel embeddings within the retained eigenspace and not in their marginal participation alone.

The bottom-tail representation showed a different organization. For *B*_2_, diagonal channel occupancy provided the strongest fixed readout, with radial terms retaining similar information and normalized angular relations being weaker. The near-full-rank regime was therefore expressed mainly through the degree to which individual measurement channels occupied the complementary bottom spectral subspace. Retained rank changed not only the magnitude of the readout but also the geometric component through which label-associated variation was expressed. The intermediate-rank and near-full-rank regimes are consequently not simply stronger and weaker versions of one common representation; they expose different spectral carriers of the integrated Pearson operator. The fully nested rank analyses provide a complementary internal check of this two-regime interpretation. Under adaptive single-rank selection, the procedure alternated primarily between *k* = 20 and *k* = 44, without converging on a single stable optimum. When two ranks were permitted, one intermediate-rank and one near-full-rank branch were retained in essentially every outer fold, with *k* = 20, 44 selected in 97 of 99 folds and *k* = 19, 44 in the remaining two. The principal value of this result lies in its representational implication, not in the higher AUC itself. Even when every rank pair was considered within the outer training data, the adaptive procedure repeatedly retained the same two parts of the spectrum highlighted by the descriptive morphology.

This pattern is consistent with the geometric decomposition showing that the two regimes are supported by different components of projector geometry: the intermediate-rank branch is predominantly relational and angular, whereas the near-full-rank branch is expressed primarily through bottom-tail occupancy. Restricting adaptive selection to a single rank can therefore discard one of two complementary spectral views of the same integrated operator.

The statistical status of this result requires an important distinction. The two-rank procedure substantially outperformed the fully nested single-rank selector, but its improvement over the fixed principal *k* = 20 representation was smaller and did not reach conventional significance. The adaptive analysis therefore does not establish that a two-rank model is superior to the fixed intermediate-rank representation. The stronger conclusion is that a fully nested procedure, with no access to the outer test subjects during rank selection, repeatedly recovered the same two spectral regimes suggested by the descriptive rank morphology and characterized by the geometric decomposition.

The decision to investigate a two-rank representation was itself motivated after inspection of the descriptive Pearson profile and is therefore exploratory. Selection of the ranks, score normalization, fusion, and evaluation of each held-out infant were nevertheless fully nested. The outer-LOSO result consequently evaluates the specified adaptive procedure without using the outer test subject for model selection, but it cannot turn a dataset-motivated hypothesis into an independently confirmed finding. Replication in an independent cohort will be needed to determine whether the same two spectral regimes and their distinct geometric carriers recur.

The rank-dependent decomposition also clarifies how projector entries should be interpreted. Off-diagonal entries of *P*_20_ are not conventional functional-connectivity edges; they are inner products between channel embeddings in the retained eigenspace. Likewise, diagonal entries of *B*_2_ are neither local activation amplitudes nor conventional node strengths. They quantify participation in the complementary bottom spectral subspace. The present findings are therefore most naturally described as rank-dependent variation in functional-network geometry, not as simple increases or decreases in pairwise connectivity.

The corresponding spatial summaries support a distributed description. Angular involvement in the intermediate-rank branch and bottom-tail occupancy in the near-full-rank branch were both spread across the measurement montage. These maps remain exploratory. Channel-level effects were not independently confirmed, and fNIRS channels sample cortical hemodynamics in measurement space and should not be treated as precise anatomical parcels. The observed patterns therefore do not establish a specific cortical circuit, a localized “bilingual” region, or a hemispheric mechanism.

The present findings are compatible with earlier analyses of the same cohort. Blanco and colleagues reported shared resting-state organization but no corrected group differences in subject-specific temporal-ICA maps or in the expression of components derived from Pearson-connectivity matrices (Blanco et al., 2021). Gao and colleagues found largely null differences in mean frontal connectivity strength together with differences in selected lateralization-related graph and directional measures (Gao et al., 2022). These studies addressed different representational targets. Group-associated variation that is weak in component amplitudes or mean connectivity can still be expressed in the geometry of a rank-constrained whole-montage subspace. The present analysis does not overturn the earlier findings; it shows that the visibility of between-group structure can depend on operator definition, temporal construction, spectral rank, spatial scope, and representation.

Pearson and coherence scores were only moderately associated, providing exploratory evidence that the two operator families preserve partly complementary subject-level geometry. Their fixed equal-weight combination increased the descriptive readout, although improvement over the stronger individual branch was borderline and both representations were characterized in the same cohort. The significance of the fusion result is consequently conceptual, not predictive: time-domain correlation and frequency-specific coherence appear to preserve overlapping but nonidentical aspects of functional-network organization. The fusion analysis should not be interpreted as an optimized multimodal classifier.

Several limitations constrain interpretation. The language groups were observational and were not experimentally assigned, so the results cannot establish a causal effect of bilingual exposure. The two monolingual subgroups were combined, leaving open whether the observed geometry is shared across Spanish- and Basque-monolingual infants. The sample was modest and group sizes were unequal, although the principal Pearson representation remained informative across repeated balanced cohorts and across positional strides from 5 to 50 s. These controls address specific sensitivity concerns but do not substitute for independent replication.

Statistical validation also differed between operator families. The fixed principal Pearson configuration underwent conditional label-permutation testing, but this analysis did not correct for the preceding descriptive rank–scale exploration. No corresponding standalone permutation analysis was performed for coherence, whose cross-fitted readouts should consequently be interpreted descriptively. More generally, cross-validation within a single cohort does not establish generalization to a new population (Varoquaux et al., 2017; Scheinost et al., 2019).

The near-full-rank decomposition and the decision to examine an adaptive two-rank representation carry an additional limitation because both were motivated after inspection of the Pearson rank profile. The matched *P*_20_–*B*_2_ decomposition provides a mechanistic interpretation of an observed secondary regime, while the nested two-rank analysis evaluates an exploratory model class and not a preregistered hypothesis. The near-uniform selection of an intermediate-rank branch together with *k* = 44 nevertheless provides a useful internal consistency result. Independent data will be required to determine whether this two-regime organization is reproducible.

Measurement limitations also remain. fNIRS captures slow hemodynamic fluctuations with limited depth sensitivity and spatial resolution, and residual systemic physiology may persist after nuisance regression (Scholkmann et al., 2014; Tachtsidis and Scholkmann, 2016). Coherence was restricted to a low-frequency band and depended on window duration and multitaper settings, while Pearson correlation summarized broadband temporal association. The operator comparison should therefore be understood as a comparison of deliberately different functional constructions, not as a controlled experiment in which the dependence estimator was the only factor that changed.

Future work should test whether the intermediate-rank regimes, operator-dependent reversal of temporal construction, secondary bottom-tail organization, and two-branch Pearson structure replicate across independent datasets, developmental stages, and imaging modalities. Direct measures of eigenspace stability, spectral gaps, and scale-to-scale subspace distances may help determine whether transitions between representational carriers correspond to identifiable changes in spectral geometry. It will also be important to separate label-free geometric reproducibility from label-associated variation and to determine which aspects of rank–scale morphology generalize beyond the language-group contrast used here as an external readout.

In summary, dominant-subspace projectors provide a basis-invariant, matrix-valued representation of functional-network geometry. In infant resting-state fNIRS, the framework revealed operator-specific rank–scale organization and an operator-dependent reversal in the order of temporal integration and spectral projection. The principal intermediate-rank Pearson regime was predominantly relational, whereas a secondary near-full-rank regime was expressed mainly through occupancy of the complementary bottom spectral subspace. A two-rank analysis, motivated post hoc but evaluated using fully nested rank selection, repeatedly retained both regimes, consistent with their carrying complementary aspects of the integrated Pearson operator. Temporal scale, retained rank, operator definition, and temporal construction jointly determine which aspects of functional-network geometry become visible.

## Acknowledgments

The authors thank the creators of the RS4 dataset for making the data and preprocessing pipeline publicly available. During the development of this work and preparation of the manuscript, the authors used ChatGPT (OpenAI) as an assistive tool for methodological discussion, code development and debugging, language refinement, manuscript organization, and text editing. All AI-assisted material and suggestions were critically evaluated and, where applicable, verified and revised by the authors. The authors take full responsibility for the study design, implementation, analyses, reported results, and interpretations.

## Supporting Information

The Supporting Information contains temporal-window sampling and robustness details, fixed-configuration statistical summaries, coherence controls and sensitivity analyses, alternative dependence estimators and operator constructions, operation-order diagnostics, nested rank-selection diagnostics, detailed geometric-decomposition results, exploratory cross-operator complementarity analyses, and additional implementation details.

## Author Contributions

**Vladi Sorkin:** Data curation, Formal analysis, Investigation, Methodology, Software, Validation, Writing – review & editing.

**Yakir Menahem:** Data curation, Investigation, Methodology, Resources, Software, Validation, Writing – review & editing.

**Dmitry Patashov:** Formal analysis, Investigation, Methodology, Validation, Writing – review & editing.

**Michal Balberg:** Formal analysis, Investigation, Methodology, Supervision, Writing – review & editing.

**Dmitry Goldstein:** Conceptualization, Formal analysis, Methodology, Software, Validation, Visualization, Writing – original draft, Writing – review & editing.

All authors reviewed and approved the final version of the manuscript.

## Data and Code Availability

The analysis-ready fNIRS data used in this study are publicly available through the Open Science Framework at https://doi.org/10.17605/OSF.IO/7FZKM (Blanco et al., 2022).

The analysis code required to reproduce the principal results reported in this article will be made available to the editors and reviewers during peer review. A curated version of the analysis code, together with environment specifications and reproducibility instructions, will be deposited in a persistent public repository upon acceptance of the manuscript. No new data were generated for this study.

