## Supporting Information for "Spectral projectors of temporally integrated functional brain networks"

#### 1 Scope of the Supporting Information

This document supplements the temporal-window robustness, fixed-configuration inference, coherence controls and sensitivity analyses, alternative dependence estimators and operator constructions, operation-order comparisons, nested rank-selection diagnostics, matched rank-dependent geometric decomposition, and exploratory cross-operator complementarity analyses reported in the main article. Temporal windows were used only to construct one representation per infant; they were not treated as independent observations.

The complete Pearson and coherence rank profiles over $k = 1, \ldots, 45$ are displayed in the main article. Rank $k = 46$ was excluded because the full-rank projector equals the identity for every subject and therefore contains no between-subject variation.

#### 2 Pearson positional-sampling sensitivity

Stride sensitivity was evaluated at the fixed Pearson configuration $w = 140$ s and $k = 20$. Because sample counts must be integral, the realized stride differed slightly from its requested duration. The centered grid was recomputed independently for each stride. Across all seven grids, the integrate-then-project AUC remained between 0.7910 and 0.8192. The 5-s and 10-s grids yielded AUCs of 0.8192 and 0.8188, respectively.

**Table S1: Centered-window sampling at $w = 140$ s.** The window length was 1,250 samples. The realized stride equals the integer sample stride divided by $f_s = 8.93$ Hz.

| Requested stride (s) | Stride (samples) | Realized stride (s) | Windows |
|---|---|---|---|
| 5 | 45 | 5.039 | 84 |
| 10 | 89 | 9.966 | 43 |
| 15 | 134 | 15.006 | 28 |
| 20 | 179 | 20.045 | 21 |
| 30 | 268 | 30.011 | 14 |
| 40 | 357 | 39.978 | 11 |
| 50 | 446 | 49.944 | 9 |

#### 3 Robustness to unequal group size

The fixed Pearson representation and readout were rerun in 50 balanced-cycle cohorts. Every cohort contained all 36 bilingual infants and 36 of the 63 monolingual infants. Monolingual inclusion was rotated as evenly as possible.

**Table S2: Balanced-cohort robustness of the fixed Pearson representation.** BA denotes balanced accuracy.

| Summary | AUC | BA | Details |
|---|---|---|---|
| Individual 72-subject cohorts | 0.7557 mean | — | 50 cohorts; median 0.7635; SD 0.0378 |
| Subject-aggregated probabilities | 0.7945 | 0.7540 | 28–29 monolingual and 50 bilingual predictions per infant |
| Equal-count aggregation | 0.7945 mean | — | 1,000 resamples; 2.5th–97.5th percentiles: 0.7884–0.8011 |

#### 4 Conditional label-permutation inference

The fixed Pearson configuration was evaluated using 9,999 global label permutations. Projector features and outer-fold partitions were fixed, while PLS regression and logistic regression were refitted inside every permuted outer training fold. This test was conditional on the specified representation and readout and did not correct for the preceding descriptive rank–scale exploration.

**Table S3: Permutation summary for the fixed Pearson configuration.**

| Quantity | Value |
|---|---|
| Observed AUC | 0.8188 |
| Number of permutations | 9,999 |
| Permuted AUC at least observed | 1 |
| One-sided Monte Carlo $p$ | 0.0002 |
| Permutation mean AUC | 0.4979 |
| Permutation SD | 0.0843 |
| Permutation 2.5th–97.5th percentiles | 0.3364–0.6631 |

#### 5 Coherence controls and taper-count sensitivity

The representative coherence analysis used the 0.03–0.08-Hz band, time–bandwidth product $NW = 3$, five DPSS tapers, $w = 260$ s, and $k = 20$. The principal representation was the mean of local rank-$k$ coherence projectors. Controls that did not preserve dominant-subspace geometry were substantially weaker.

Taper-count sensitivity was evaluated at the fixed representative configuration while holding the frequency band, $w = 260$ s, $k = 20$, and $NW = 3$ unchanged. Performance was weaker with

**Table S4: Coherence controls and taper-count sensitivity.** Representation and control summaries. Values marked as approximate are rounded summaries from the exploratory control analyses.

| Representation or control | Readout | Interpretation |
|---|---|---|
| Mean of local projectors, representative (260 s, $k = 20$) | AUC 0.8126; BA 0.7004 | Fixed representative configuration |
| Mean of local projectors, largest descriptive grid cell (270 s, $k = 21$) | AUC 0.8320 | Descriptive maximum, not a validated optimum |
| Position-averaged raw coherence | Best AUC approximately 0.62 | Substantially weaker control |
| Channel-wise log band power | AUC approximately 0.60–0.61 | Substantially weaker control |

two tapers and remained informative across $K = 3$–5.

**Table S4 (continued): Coherence controls and taper-count sensitivity.** Fixed-configuration taper-count readouts at $w = 260$ s and $k = 20$.

| DPSS tapers $K$ | AUC | BA | Specificity | Sensitivity |
|---|---|---|---|---|
| 2 | 0.7090 | 0.5774 | 0.6825 | 0.4722 |
| 3 | 0.7884 | 0.7004 | 0.7619 | 0.6389 |
| 4 | 0.8020 | 0.7123 | 0.7302 | 0.6944 |
| 5 | 0.8126 | 0.7004 | 0.7619 | 0.6389 |

Using $K = 5$ as the reference at the fixed representative configuration, the paired bootstrap AUC difference relative to $K = 2$ was 0.1036 (95% interval 0.0018–0.2108; two-sided $p = 0.0468$). Differences relative to $K = 3$ and $K = 4$ were smaller and their intervals included zero: $\Delta$AUC = 0.0243 (95% interval −0.0344–0.0877; $p = 0.4335$) and $\Delta$AUC = 0.0106 (95% interval −0.0278–0.0511; $p = 0.5823$), respectively. Thus, at the representative 260-s scale, the coherence readout was not dependent on a single taper count within the $K = 3$–5 range.

To determine whether this conclusion was specific to the representative window, we additionally compared $K = 3, 4, 5$ across neighboring window durations of 250–280 s while holding the retained rank fixed at $k = 20$. This was a temporal-scale sensitivity analysis at a fixed rank, not an additional rank-selection analysis.

**Table S4 (continued): Coherence controls and taper-count sensitivity.** Focused taper-by-window sensitivity at fixed $k = 20$. Entries are outer leave-one-subject-out AUC, with balanced accuracy in parentheses.

| Window (s) | $K = 3$ | $K = 4$ | $K = 5$ |
|---|---|---|---|
| 250 | 0.7760 (0.6845) | 0.7593 (0.6528) | 0.8135 (0.7202) |
| 260 | 0.7884 (0.7004) | 0.8020 (0.7123) | 0.8126 (0.7004) |
| 270 | 0.7637 (0.6925) | 0.7571 (0.7063) | 0.8153 (0.7361) |
| 280 | 0.7703 (0.6667) | 0.7813 (0.6845) | 0.7998 (0.7004) |
| Mean AUC across windows | $0.7746 \pm 0.0105$ | $0.7749 \pm 0.0211$ | $0.8103 \pm 0.0071$ |

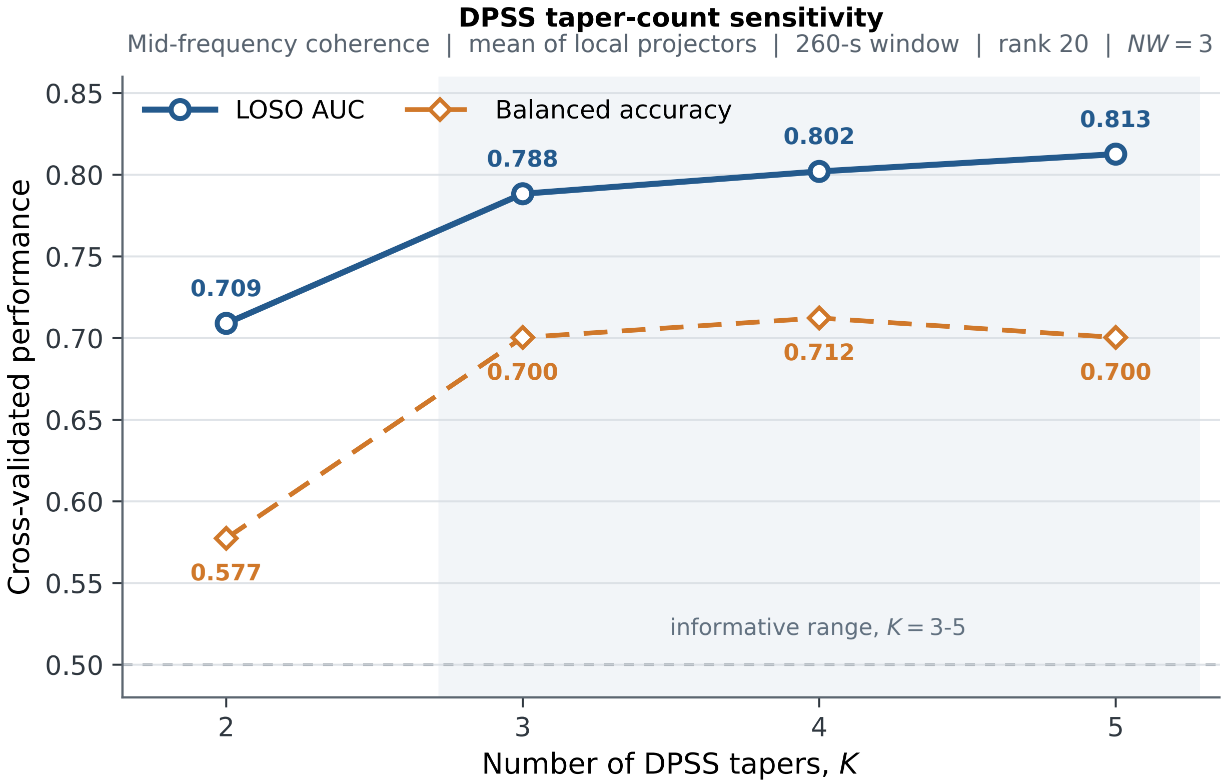


**Figure S3: Sensitivity of the representative coherence readout to DPSS taper count.** Outer leave-one-subject-out AUC and balanced accuracy for the project–integrate coherence representation at $w = 260$ s, $k = 20$, and $NW = 3$, using $K = 2, 3, 4, 5$ DPSS tapers. Performance was weaker for $K = 2$ and remained informative across $K = 3$–$5$.

The five-taper representation yielded the highest AUC at each of the four neighboring window durations. Paired bootstrap differences between $K = 5$ and $K = 3$ did not exclude zero at any of the four windows. For $K = 5$ versus $K = 4$, the 95% interval excluded zero at 250 s ($\Delta$AUC = 0.0542, interval 0.0101–0.1032, $p = 0.0152$) and 270 s ($\Delta$AUC = 0.0582, interval 0.0084–0.1142, $p = 0.0252$), but not at 260 or 280 s. The neighboring-window analysis therefore supports persistence of the coherence readout across moderate taper counts while also showing that $K = 5$ provided the most consistently high AUC across this local temporal range.

### 6 Robustness to dependence estimation and operator construction

Focused exploratory analyses assessed whether rank-dependent organization was specific to Pearson correlation or to the construction of the underlying operator. Pearson-style integration-before-projection was repeated using Spearman and Gaussian-copula dependence estimates over window durations of 120–160 s and ranks $k = 1, \ldots, 45$. Both alternatives retained an elevated intermediate-rank regime. A separate heat-kernel graph construction produced distinct low- and high-rank regimes. These analyses were interpreted from the complete profiles rather than from isolated numerical maxima.

**a**

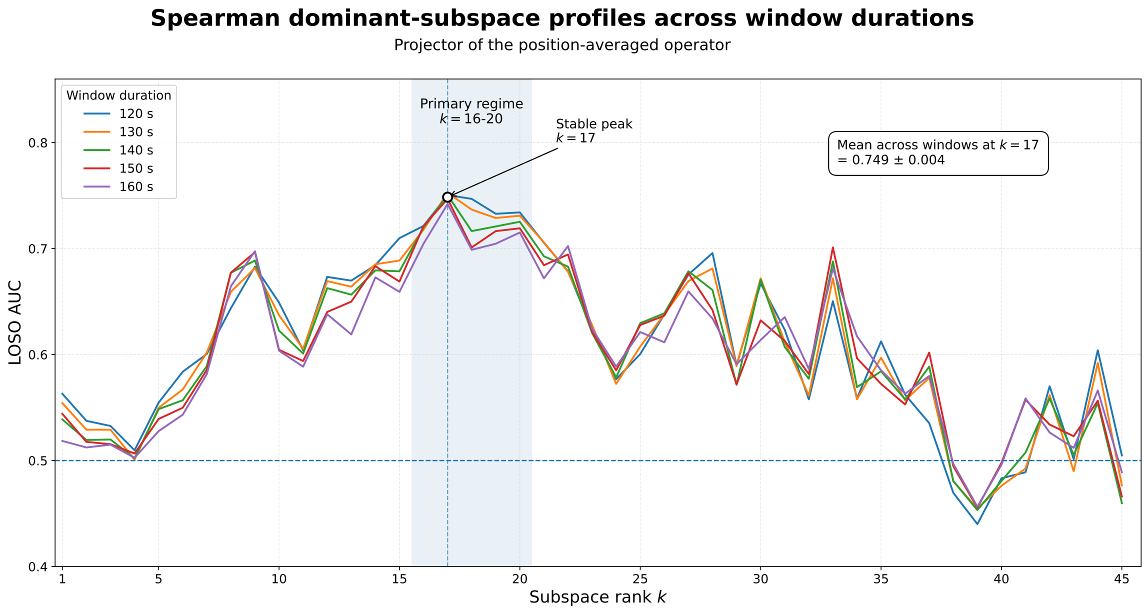


**b**

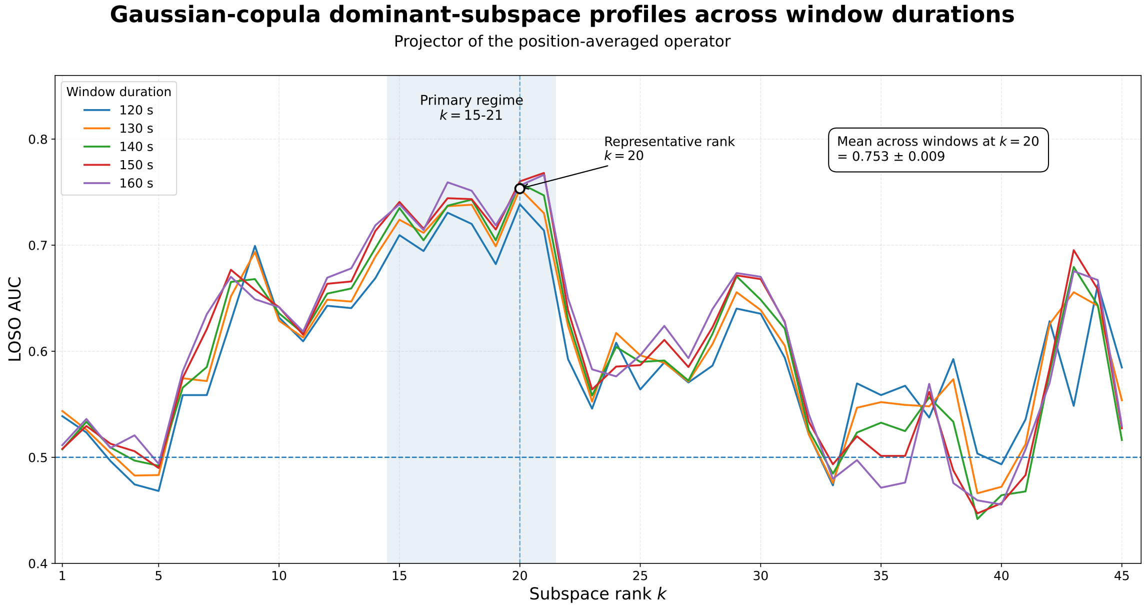


**Figure S4: Sensitivity to the dependence estimator.** **(a)** Spearman dependence produced an elevated regime over approximately $k = 16$–$20$, with the largest across-window mean at $k = 17$ ($\mathrm{AUC} = 0.749 \pm 0.004$). **(b)** Gaussian-copula dependence produced a similar regime over approximately $k = 15$–$21$; the representative rank $k = 20$ yielded $\mathrm{AUC} = 0.753 \pm 0.009$. Values following $\pm$ are standard deviations across the five window durations. Outer leave-one-subject-out AUC is shown for the complete range $k = 1, \ldots, 45$. These profiles were exploratory robustness checks and isolated maxima were not treated as validated optima.

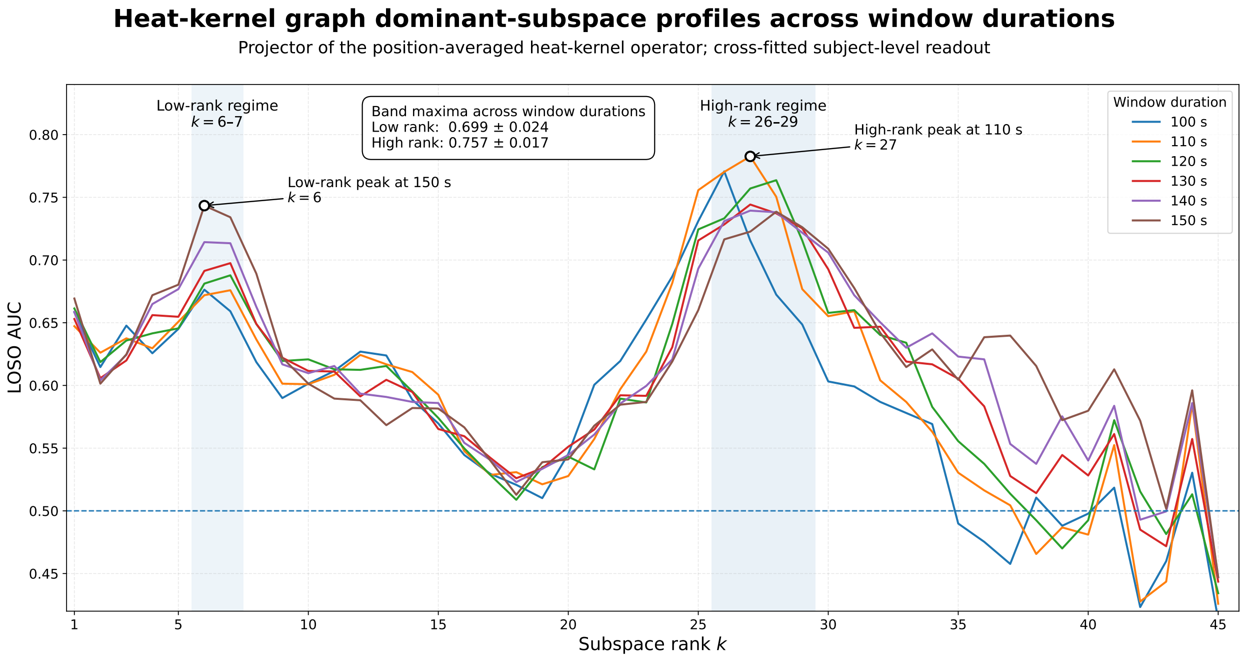


**Figure S5: Sensitivity to heat-kernel graph construction.** Outer leave-one-subject-out AUC for the heat-kernel graph dominant-subspace representation across $k = 1, \ldots, 45$ and window durations of 100–150 s. Two distinct elevated regimes were observed, at $k = 6$–7 and $k = 26$–29. Across window durations, the within-band maximum AUC was $0.699 \pm 0.024$ for the low-rank regime and $0.757 \pm 0.017$ for the high-rank regime (mean $\pm$ SD across the six window durations). Rank-dependent organization therefore persisted under smooth graph-kernel construction, although its precise spectral location differed from that of the correlation-based operators. This analysis was exploratory.

### 7 Temporal operation order

The two primary operator families showed opposite preferences for the order of temporal integration and spectral projection. For Pearson, projecting the position-averaged operator was favored. For coherence, averaging locally estimated projectors was favored.

For clarity, let

$$\overline{C} = \frac{1}{H}\sum_{h=1}^{H} C_h$$

denote the temporally integrated operator, and let

$$\overline{P_k(C)} = \frac{1}{H}\sum_{h=1}^{H} P_k(C_h)$$

denote the mean of the local rank-$k$ projectors. Thus, $P_k(\overline{C})$ represents integration before projection, whereas $\overline{P_k(C)}$ represents projection before integration.

**Table S5: Fixed-configuration operation-order comparisons.** The paired difference is oriented toward the favored construction in each operator family. Bootstrap intervals are paired 95% percentile intervals.

| Operator and construction | AUC | BA | Paired ΔAUC | 95% interval |
|---|---|---|---|---|
| Pearson: integrate then project, $Q = P_k(\overline{C})$ | 0.8188 | 0.7698 | 0.2218 | 0.1234–0.3254 |
| Pearson: project then integrate, $A = \overline{P_k(C)}$ | 0.5970 | 0.6052 | — | — |
| Pearson: reprojected occupancy, $P_k(A)$ | 0.5670 | — | 0.0300[a] | −0.0573–0.1169 |
| Coherence: project then integrate, $A = \overline{P_k(C)}$ | 0.8126 | 0.7004 | 0.1124 | 0.0322–0.1953 |
| Coherence: integrate then project, $Q = P_k(\overline{C})$ | 0.7002 | 0.6488 | — | — |

[a]Unreprojected occupancy minus reprojected occupancy.

At the largest descriptive coherence grid cell, $w = 270$ s and $k = 21$, the corresponding AUCs were 0.8320 for projection before integration and 0.6539 for integration before projection.

**Table S6: Label-free Pearson diagnostics at $w = 140$ s and $k = 20$.** Here $Q = P_k(\overline{C})$ is the projector of the integrated operator and $A = \overline{P_k(C)}$ is the mean-local-projector matrix.

| Diagnostic | Mean across subjects |
|---|---|
| Frobenius idempotence error, $\|A^2 - A\|_F$ | 1.090 |
| Normalized Frobenius distance, $\|Q - A\|_F/\sqrt{2k}$ | 0.322 |
| Normalized overlap, $\mathrm{tr}(QA)/k$ | 0.734 |

#### 8 Matched geometric decomposition of the Pearson rank regimes

The principal intermediate-rank projector $P_{20}$ and the secondary near-full-rank representation were decomposed using the same five feature families. For the latter, the invariant complementary projector

$$B_2 = I - P_{44}$$

was analyzed rather than individual bottom eigenvectors. In the 46-channel system, $B_2$ spans the two-dimensional bottom spectral subspace and contains the same subspace information as $P_{44}$.

**Table S7: Matched geometric decomposition of the intermediate- and near-full-rank Pearson regimes.** All feature families were evaluated with the same strict outer-LOSO readout.

| Representation | Feature family | Coordinates | AUC |
|---|---|---:|---:|
| $P_{20}$ | Diagonal channel leverage | 46 | 0.490 |
| $P_{20}$ | Pairwise radial terms | 1,035 | 0.495 |
| $P_{20}$ | Normalized angular relations | 1,035 | 0.771 |
| $P_{20}$ | Raw off-diagonal entries | 1,035 | 0.783 |
| $P_{20}$ | Complete matrix | 1,081 | 0.819 |
| $B_2 = I - P_{44}$ | Diagonal bottom-tail occupancy | 46 | 0.799 |
| $B_2 = I - P_{44}$ | Pairwise radial terms | 1,035 | 0.784 |
| $B_2 = I - P_{44}$ | Normalized angular relations | 1,035 | 0.698 |
| $B_2 = I - P_{44}$ | Raw off-diagonal entries | 1,035 | 0.759 |
| $B_2 = I - P_{44}$ | Complete matrix | 1,081 | 0.785 |

The carrier pattern changed qualitatively with rank. For $P_{20}$, diagonal leverage and radial terms were essentially non-informative in isolation, whereas normalized angular relations and raw off-diagonal entries retained most of the complete-projector readout. The principal intermediate-rank regime was therefore predominantly relational. For $B_2$, diagonal bottom-tail occupancy provided the strongest fixed readout, with radial terms retaining similar information and angular relations being weaker. The near-full-rank regime was therefore expressed primarily through channel-wise occupancy of the complementary lower spectral subspace. Individual channel and pairwise spatial effects remain exploratory measurement-space summaries.

#### 9 Fully nested adaptive rank-selection diagnostics

Adaptive rank selection was evaluated at the fixed Pearson temporal scale $w = 140$ s over all candidate ranks $k = 1, \ldots, 45$. For every outer leave-one-subject-out fold, candidate ranks were compared only within the outer training set using a common stratified nine-fold inner cross-validation. The two-rank analysis was motivated after inspection of the descriptive rank profile, but rank-pair selection, score standardization, refitting, and evaluation of the held-out infant were fully nested.

The two-rank readout exceeded nested single-rank selection by $\Delta\text{AUC} = 0.1301$ (95% paired-bootstrap interval 0.0551–0.2156; two-sided $p = 0.0004$). Relative to the fixed $k = 20$ representation,

**Table S8: Adaptive Pearson rank-selection readouts.** The fixed $k = 20$ result is included as a reference and was not selected within the nested procedures.

| Readout | AUC | BA | Comparison |
|---|---|---|---|
| Fixed $k = 20$ representation | 0.8188 | 0.7698 | Reference principal representation |
| Fully nested single-rank selection | 0.7465 | 0.7123 | $\Delta$AUC = −0.0723 versus fixed $k = 20$; 95% interval −0.1455–−0.0057 |
| Fully nested two-rank equal fusion | 0.8765 | 0.7917 | 95% AUC interval 0.7862–0.9506 |

the increase was smaller, $\Delta\text{AUC} = 0.0578$, and the paired interval included zero (−0.0018–0.1235; $p = 0.0594$). The adaptive result is therefore interpreted primarily as evidence that the two rank regimes carry complementary information, not as a conventionally significant improvement over the fixed principal representation.

**Table S9: Rank-selection frequencies across outer LOSO folds.**

| Procedure | Selected rank or pair | Outer folds | Fraction |
|---|---|---|---|
| Single-rank | $k = 20$ | 72 | 72.7% |
| Single-rank | $k = 44$ | 25 | 25.3% |
| Single-rank | $k = 14$ | 1 | 1.0% |
| Single-rank | $k = 17$ | 1 | 1.0% |
| Two-rank | $(k_a, k_b) = (20, 44)$ | 97 | 98.0% |
| Two-rank | $(k_a, k_b) = (19, 44)$ | 2 | 2.0% |

The unrestricted single-rank selector alternated mainly between the intermediate-rank and near-full-rank regimes. In contrast, the two-rank procedure retained $k = 44$ in every outer fold and an intermediate-rank branch in every fold. Because pair optimization was performed using pooled inner out-of-fold AUC within each outer training set, the concentration of the selected pair provides a direct diagnostic of selection stability. It does not make the two-rank hypothesis confirmatory: the decision to allow two branches was motivated after inspection of the descriptive rank morphology.

#### 10 Exploratory cross-operator complementarity

Pearson and coherence branch scores were only moderately associated (Pearson $r = 0.337$, $p = 0.00064$; Spearman $\rho = 0.307$, $p = 0.0020$). Their binary predictions differed for 41.4% of infants.

The fusion improvement over coherence was 0.0785 (95% interval 0.0194–0.1429; $p = 0.0084$); the improvement over Pearson was 0.0723 (95% interval −0.0013–0.1477; $p = 0.0540$). Comparison with the better-performing branch within each bootstrap sample was borderline ($p = 0.0624$). A post hoc fixed-weight sweep improved AUC by only 0.0022 beyond the equal-weight combination. The fusion is therefore descriptive and exploratory rather than a separately optimized predictive model.

**Table S10: Branch and equal-weight fusion readouts.** Fusion used an untrained equal-weight mean of the two saved outer-LOSO logits.

| Readout | AUC | BA |
|---|---|---|
| Pearson | 0.8188 | 0.7698 |
| Coherence | 0.8126 | 0.7004 |
| Equal-weight logit fusion | 0.8911 | 0.8175 |

For completeness, both branches were correct for 53 infants, Pearson alone was correct for 23, coherence alone for 18, and both were incorrect for 5.

#### 11 Implementation details

All matrices were represented in the original 46-channel coordinates. Projector and occupancy matrices used the upper triangle including the diagonal (1,081 coordinates); baselines with invariant unit diagonal used the strict upper triangle (1,035 coordinates). Within each outer LOSO training fold, nonfinite or effectively constant coordinates were removed, remaining features were standardized, five PLS components were estimated, and an L2-regularized class-weighted logistic regression model with $C = 0.5$ was fitted. No group labels entered window definition, operator estimation, eigenspace construction, or projector formation.

Bootstrap comparisons used 5,000 paired stratified resamples unless stated otherwise. The matched rank-dependent decomposition and nested rank-pair analyses used 10,000 bootstrap replicates. Taper-count AUC comparisons used 5,000 paired bootstrap resamples. The same resampled subjects were used for all representations in a paired comparison. The infant, not the temporal window, was the statistical unit throughout.
